# Quercetin and 6-Br-quercetin: antioxidant properties and off target screening results advance glycosylated 6BrQ as developmental candidates for Alzheimer’s disease

**DOI:** 10.64898/2026.03.10.710855

**Authors:** George R Uhl, Balaji Kannan, Emily Hess, Henderson, Kevin Schultz

## Abstract

Quercetin is an abundant dietary flavonol with interesting *in vitro* properties that include substrate-selective positive allosteric modulation (PAM) of the activity of the receptor type protein tyrosine phosphatase D (PTPRD) and substantial antioxidant actions. Its *in vivo* activities include reducing incidence of Alzheimer’s disease (AD) and reducing AD neurofibrillary pathology in mouse models. Structure-activity studies have identified quercetin analogs with improved *in vitro* and *in vivo* properties, including the improved PTPRD PAM 6-bromoquercetin (6BrQ). However, there is no comparison of the antioxidant properties of 6BrQ to those of quercetin. There is no systematic screening for activities of quercetin or of 6BrQ using a panel of targets for most currently-used drugs.

We now report that both quercetin and 6BrQ provide equivalent results in cyclic voltammetric and biochemical antioxidant assays. We also report that neither 10^-7^ M quercetin nor 6BrQ provides any significant (>50%) effects on any of the 104 assays in a Eurofins off-target screening panel. At 10^-5^ M, both quercetin and 6BrQ exert significant effects in assays for glycogen synthase kinase 3 (GSK3β) as well as those for serotonin 5HT2B receptor, adenosine transport, adenosine A2A receptors, cyclooxygenases COX1 and COX2, phosphodiesterases PDE3A and 4D2 and PPAR gamma. These data extend prior characterization of quercetin’s biochemical effects, provide novel results for 6BrQ and support the likelihood that both quercetin and 6BrQ can a) directly inhibit GSK3, b) reduce GSK3 activities *via* enhancement of its dephosphorylation by PTPRD and c) display modest numbers of off target activities at high concentrations, several of which could conceivably contribute to anti-AD activities. These results advance bioavailable glycosylated prodrugs that can be metabolized to 6BrQ as developmental candidates for AD.

## Background

Quercetin is one of the most abundant dietary flavonols (1). Higher intake of quercetin and other flavonols reduces incidence of Alzheimer’s disease (AD), especially in individuals who have increased genetic risks (2). We have identified quercetin as a lead compound positive allosteric modulator of the receptor type protein tyrosine phosphatase D (PTPRD)’s ability to dephosphorylate and downregulate activities of glycogen synthase kinase 3 α and β (3).

Quercetin’s antioxidant and other properties have been long studied. Structure-activity studies have identified quercetin analogs that display improvements in these antioxidant (4) and its other properties. We have recently completed structure-activity studies with almost 110 natural product and synthetic quercetin analogs (*data not shown*). 6-Bromo quercetin (6BrQ) provides the most robust improvement in quercetin’s PTPRD PAM activities *in vitro.* 6BrQ dramatically improves quercetin’s ability to reduce development of AD-like hyperphosphorylated tau in a mouse model for AD (5). Although quercetin is poorly bioavailable (2), better-adsorbed glycosylated prodrugs that are metabolized to 6BrQ are thus developmental candidate drugs to reduce AD neurofibrillary pathology.

Quercetin has a substantial tolerability in humans and experimental animals, leading to its generally recognized as safe (GRN # 341) designation by the Food and Drug Administration. 6BrQ has not exerted any notable toxicities in studies of wildtype C57 or Alzheimer’s disease model mice treated from weaning to 4 months of age (5).

Alternate day intraperitoneal doses in these mice were 25 mg/kg for 10 – 12 weeks. We have been unable to find data that compares antioxidant properties of 6BrQ to those of quercetin. There are no systematic screens for quercetin activity at the targets for marketed drugs that allow comparisons between this GRAS compound and our new developmental candidate active metabolite, 6BrQ.

We now thus report results of cyclic voltametric and biochemical tests for antioxidant properties of quercetin and 6BrQ. We report activities of 100nM and 10 μM concentrations of quercetin and 6BrQ in Eurofins screening assays for activities at 104 sites that include G protein coupled receptors, ligand gated and other ion channels, enzymes and transporters. These assays correspond to the targets for most currently-available drugs. The quercetin results support several observations in the literature. Taken together with biochemical and *in vivo* data, these antioxidant and off target results support development of 6BrQ as a novel agent to enhance PTPRD PAM activity while retaining quercetin’s antioxidant properties.

## Methods

### Quercetin (Sigma) was > 97% pure

**6BrQ** (6 bromo quercetin) was synthesized, purified and characterized. N-bromosuccinimide (0.605 g, 3.4 mmol) was dissolved in deionized water (200 mL) containing 6 mL of 3 M NaOH. This solution was added dropwise to quercetin (1.0 g, 3.31 mmol) in methanol (200 mL) under nitrogen. After 8 h of stirring at room temperature, sodium hydrosulfite (2.0 g, 11.5 mmol in 100 mL deionized water) was added. The reaction mixture was acidified using 3 M HCl and cooled overnight. The resulting yellow precipitate was collected by filtration and dried to give a crude product (0.807 g, 68% yield). 400 mg of the crude product was purified by flash silica column chromatography using a gradient of 70% to 90% diethyl ether/hexanes (R_f_ = 0.40 (90% diethyl ether/hexanes)) to afford a yellow powder (0.168 g, 42% yield). 300 mHz NMR verification of structure and purity came from 1H NMR in d6-acetone δ = 6.75 (s, 1H), 7.00 (d, J = 8.4 Hz, 1H), 7.70 (dd, J = 8.4, 2.1 Hz, 1H), 7.83 (d, J = 2.1 Hz, 1H), 13.05 (s, 1H), 1 H NMR in DMSO δ = 6.62 (s, 1H), 6.89 (d, J = 8.4 Hz, 1H), 7.56 (dd, J = 8.4, 2.1 Hz, 1H), 7.68 (d, J = 2.1 Hz, 1H), 9.37 (s, 1H), 9.59 (s, 1H), 9.67 (s, 1H), 11.70 (s, 1H), 13.40 (s, 1H) and 13 C NMR in d-acetone δ = 92.94, 94.77, 104.42, 115.80, 116.21, 121.49, 123.52, 136.73, 145.67, 147.60, 148.52, 156.03, 158.78, 160.91, 176.05.

### Assays used

*Cyclic voltammetry* Cyclic voltammograms of quercetin and 6BrQ (800 μM) were obtained at 22 ° C in a 15 mL three electrode cell with a 3.0 mm diameter glassy carbon working electrode that was carefully polished before each experiment with 0.05 μm alumina paste, a platinum rod counter electrode, a Ag/AgCl reference electrode, supporting electrolyte containing ethanol and K_2_HPO_4_/ KH_2_PO_4_ (both 0.05 mol/L) 1:1 (v/v) with pH adjusted with KOH or H_3_PO_4_ to 7.4 and a PAR model 263A potentiostat/galvanostat with PowerSuite software as described (6).

*Biochemical antioxidant assay* DPPH radical scavenging assays were performed as described (7) (8)(9). Methanol stock solutions contained 69 µM DPPH and 3.3 uM to 3.3 mM quercetin or BrQ. 3.0 mL cuvettes with 2.7 mL of 69 µM DPPH and the indicated final concentration of quercetin or 6BRQ were shaken vigorously and incubated in the dark at 22° C for 30 minutes. Absorbance at ∼516 nm was noted and the DPPH scavenging effect was calculated using the following equation.

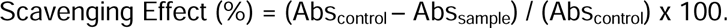

#### EUROFINS assays

Assays were carried out at EUROFINS using the methods shown in Table 1 and 10^-5^ and 10^-7^ M concentrations of quercetin and 6BrQ. “Activity” was defined as producing > 50% effects in the target assay, per EUROFINS definitions.

**Table I:**
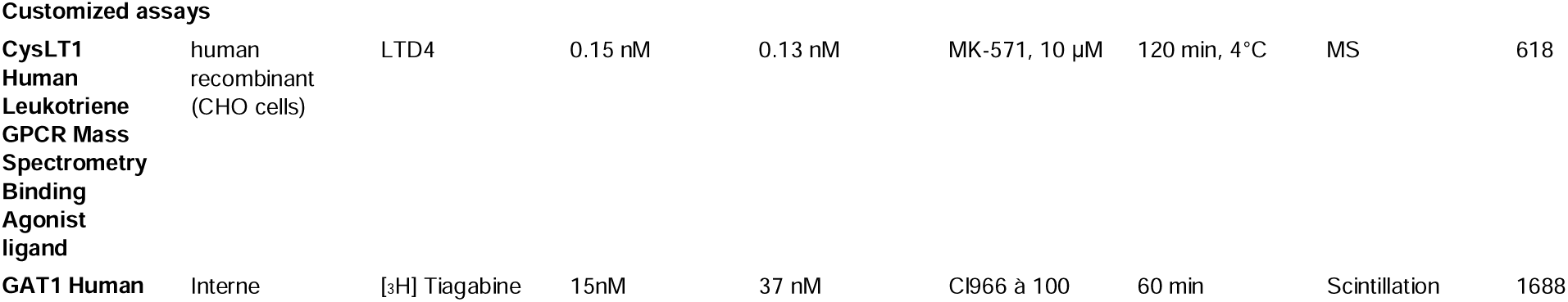
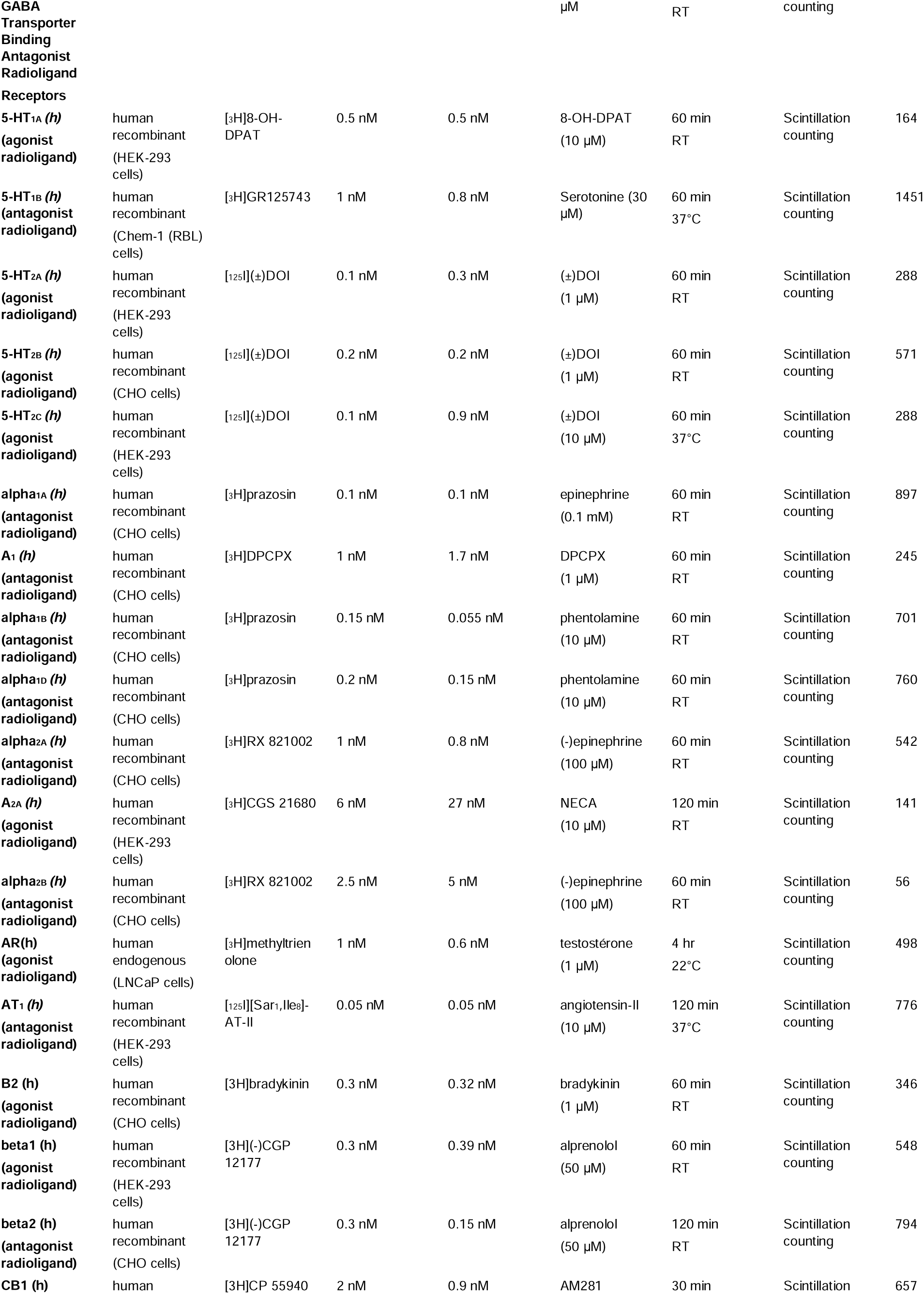
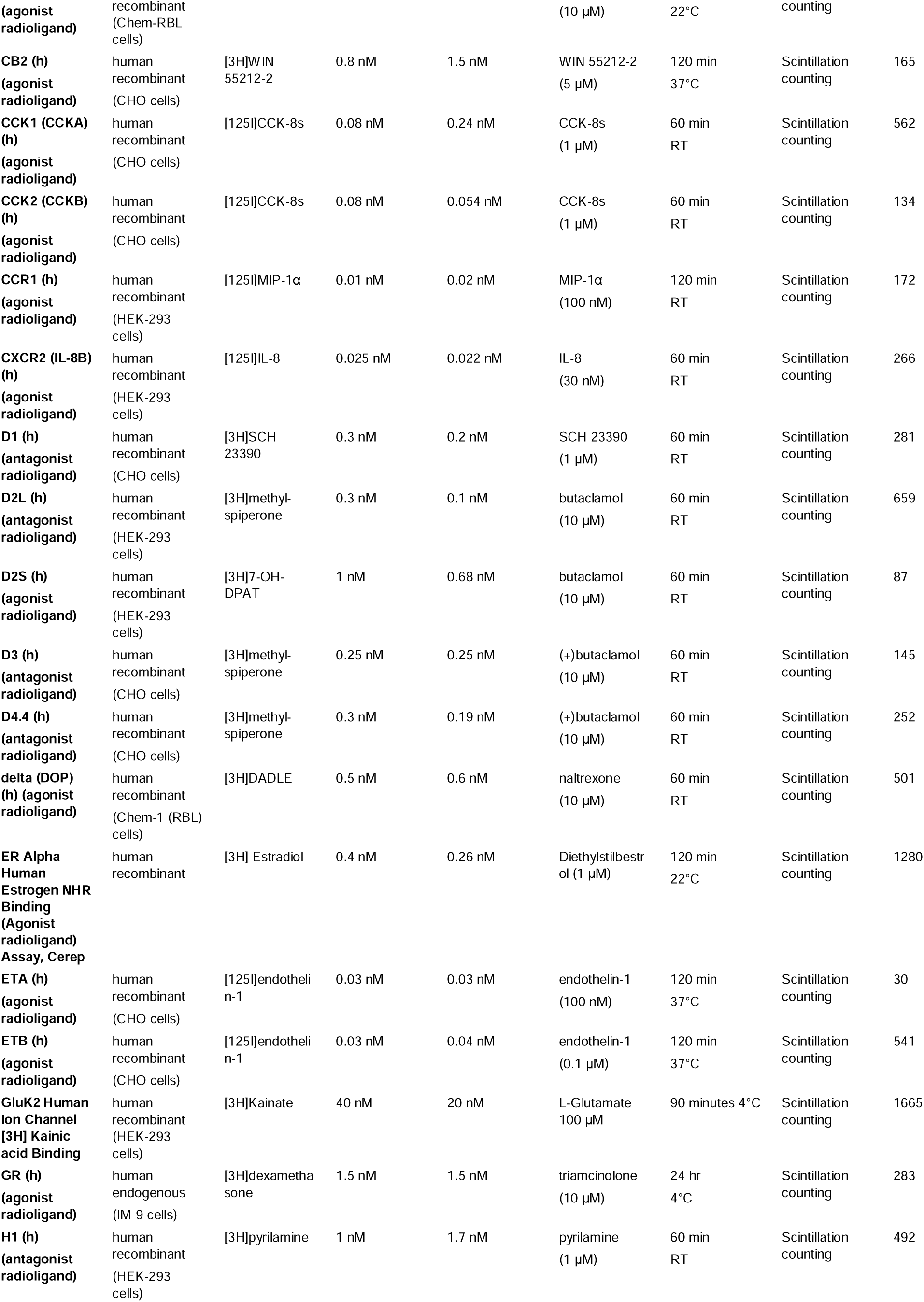
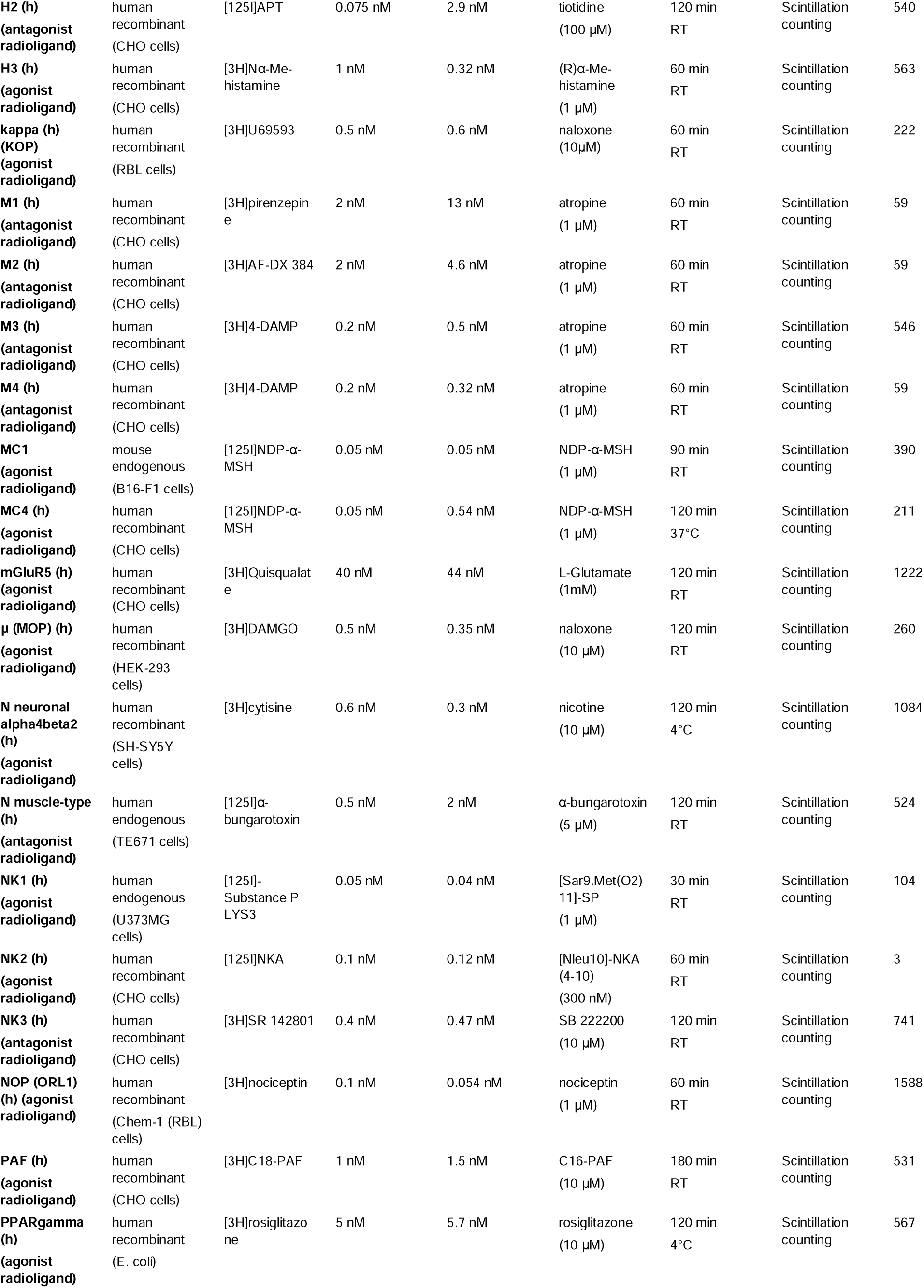
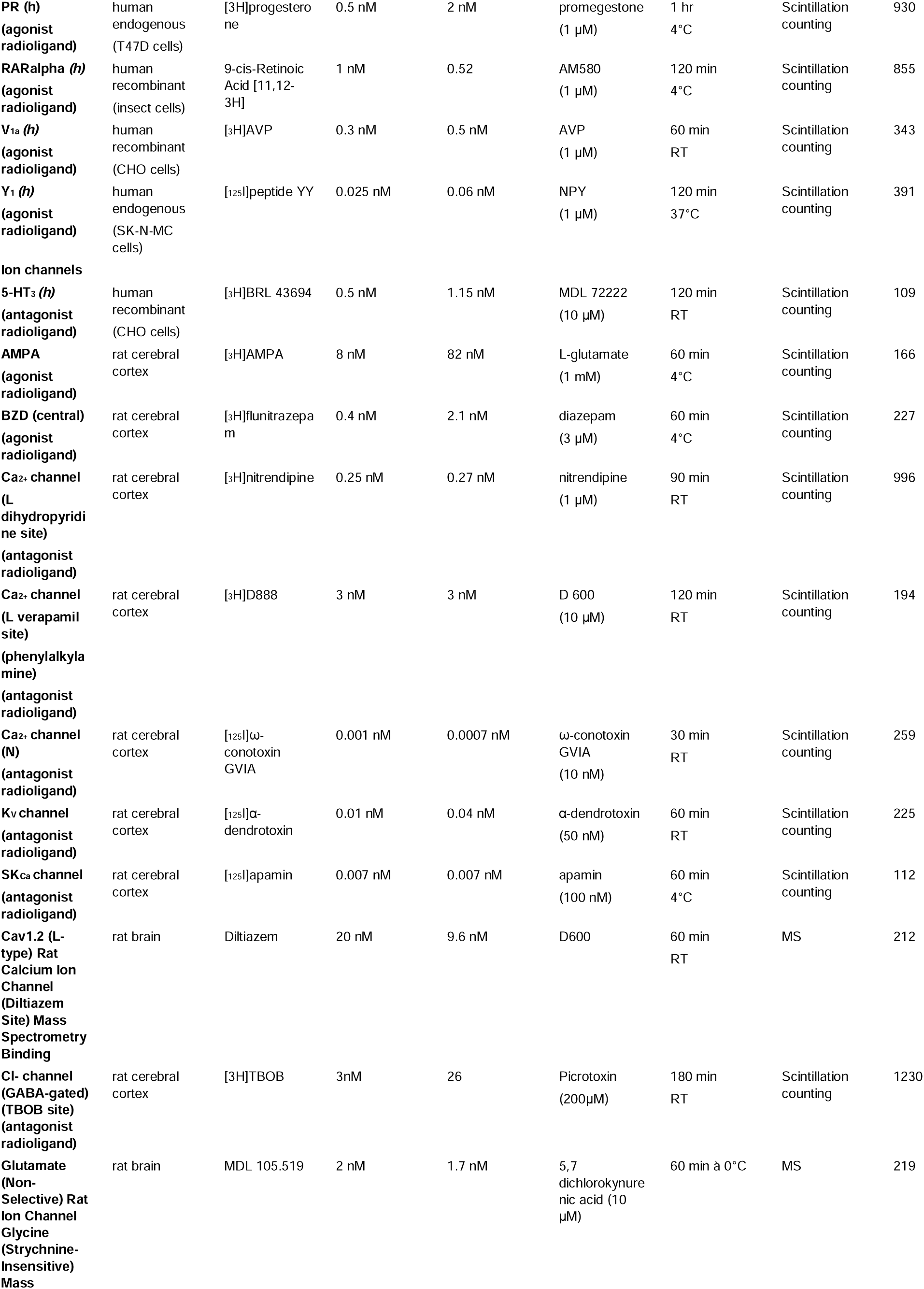
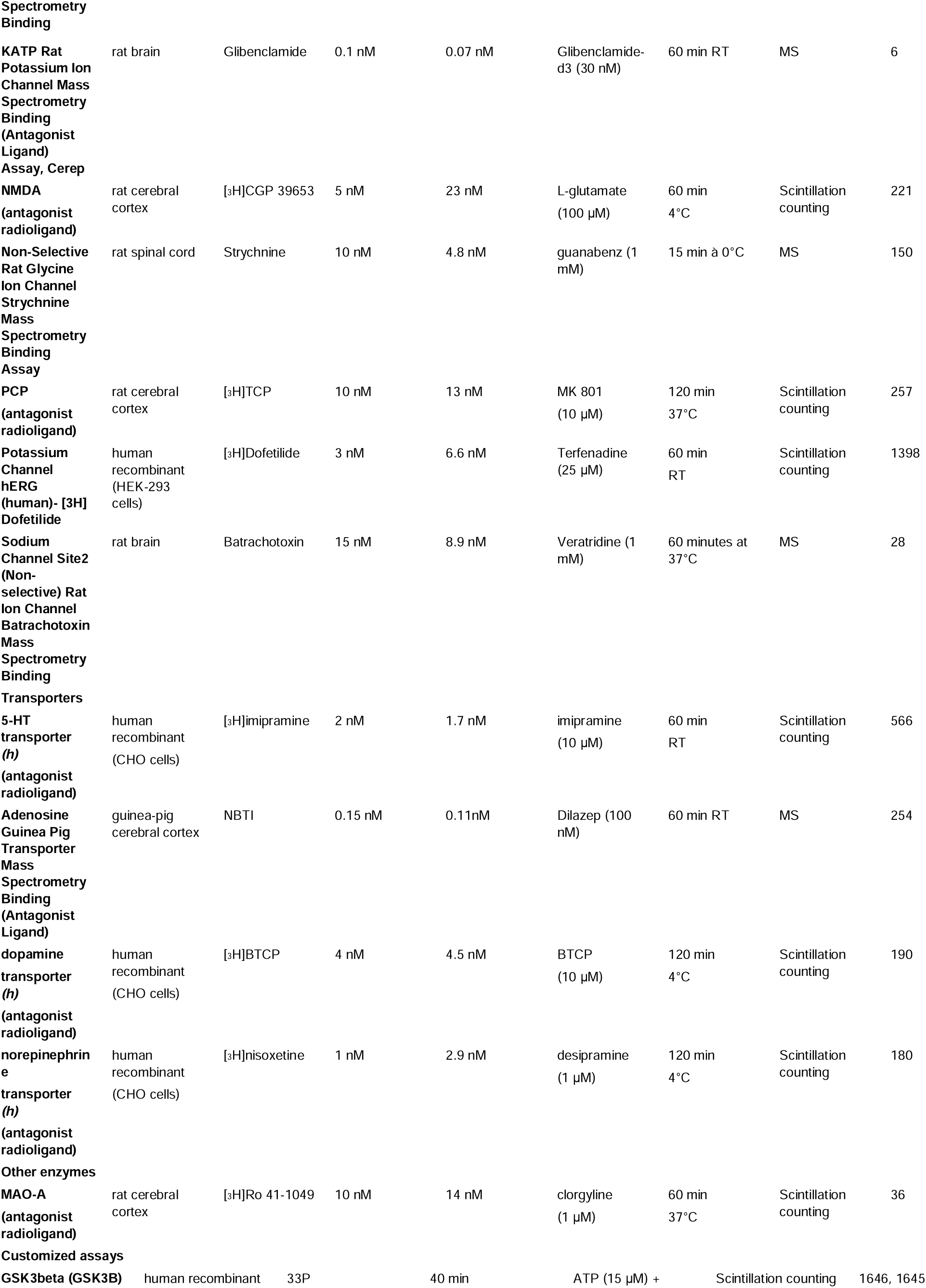
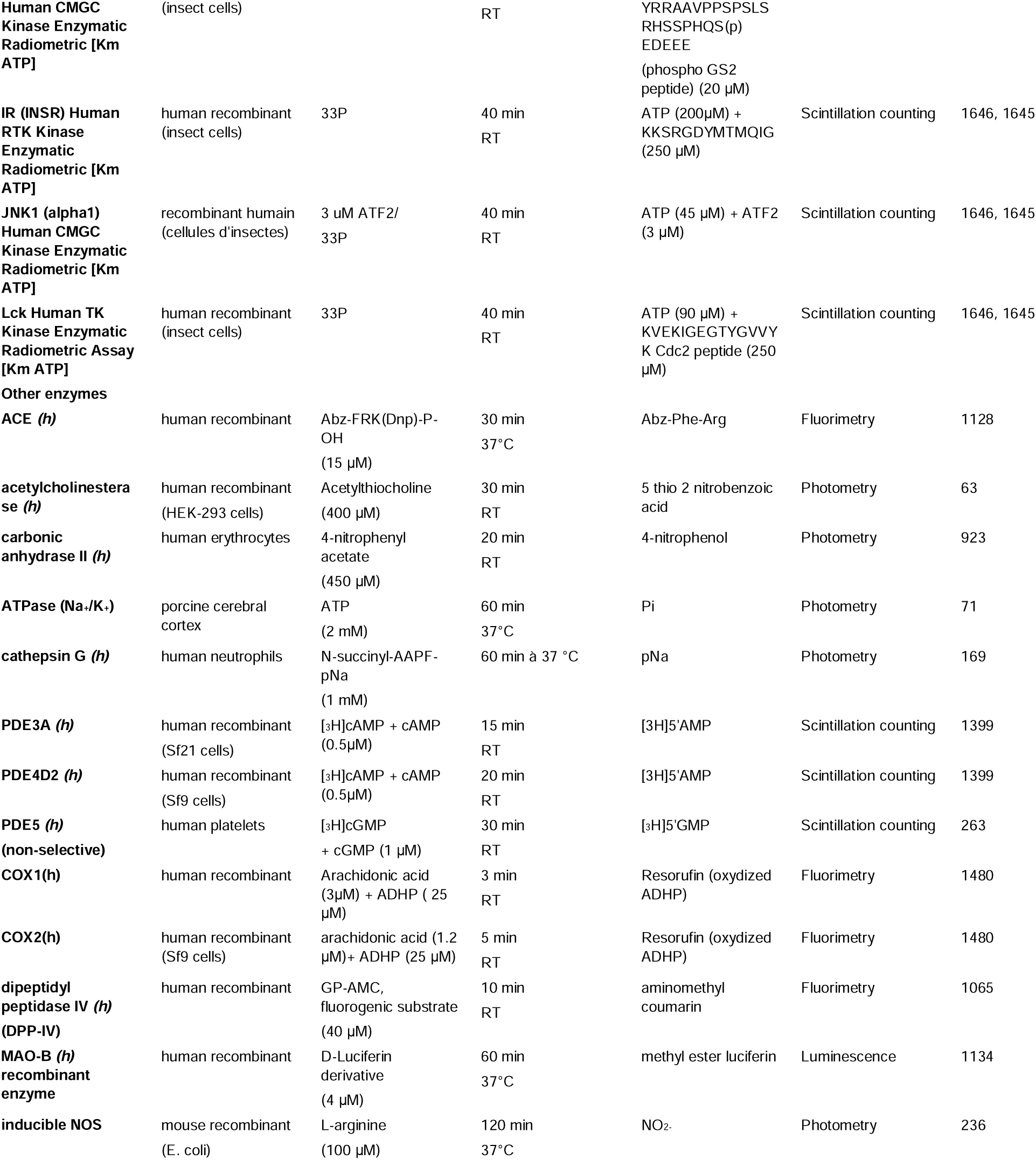
Eurofins assays used herein.

## Results

Both quercetin and 6BrQ displayed good antioxidant properties in both cyclic voltametric and biochemical antioxidant assays (Figs 1,2). 6BrQ retains the full antioxidant properties of quercetin in each of these assays.

**Fig 1:**
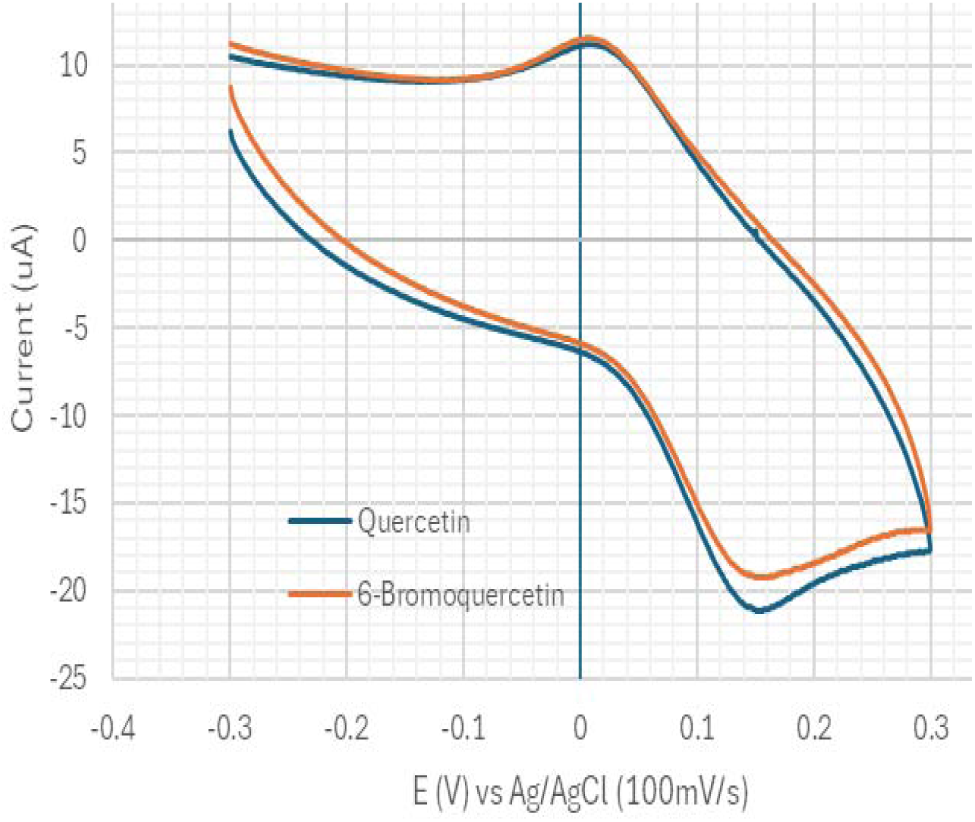
Similar cyclic voltametric resuquercetin and 6BrQ (10^-4^ M)

**Fig 2:**
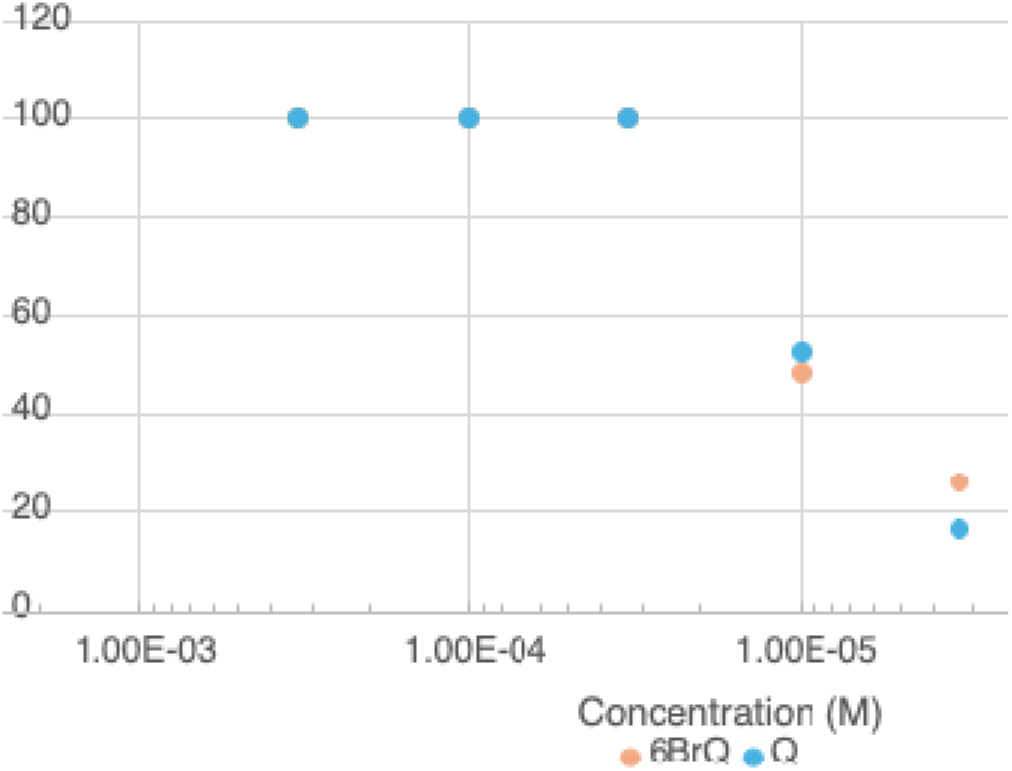
DNPPH scavenging antioxidant effect of quercetin and 6BrQ.

In EUROFINS assays, there was no activity of 10^-7^ M concentrations of either quercetin or 6BrQ on any of the receptor, ion channel, transporter or enzymatic targets. By contrast, positive controls for each of these assays were indeed positive, consistent with well-performed assays and appropriate biological activities of the receptor, channel, transporter or enzyme tested.

There were 9 Eurofins assays in which 10^-5^ M concentrations of both quercetin and 6BrQ produced >50% effects (Table II).

**Table II:**
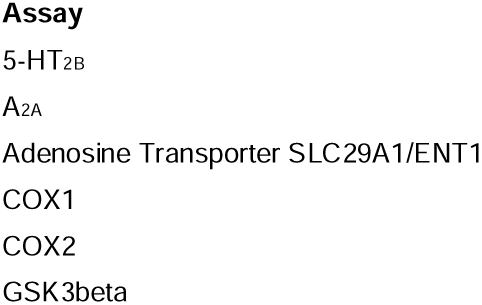
Assays in which both quercetin and 6BrQ provide > 50% effects at 10-5 M.

The only assay in which 6BrQ provided a > 50% effect that was not provided by quercetin was for CCK1 receptor activity. By contrast, quercetin provided activities at A1, D4.4, ER Alpha, MAO-A, PDE5 and PR that were not seen with 6BrQ.

## Discussion

Interest in quercetin and motivation for structure-activity relationship studies based on quercetin have come from quercetin *in vitro* and *in vivo* activities, even when all of these activities have not been fully characterized. For example, our interest in quercetin and its analogs came from our initially-surprisingly observations that quercetin provided robust positive allosteric modulation of the ability of PTPRD’s phosphatase to release phosphate from regulatory phosphotyrosines of GSK3 α and β (3). GSK3s are prominent contributors to the pathological hyperphosphorylation of tau in AD.

Others have found that a) higher dietary intake of quercetin and other flavonols was associated with strikingly lower incidence of Alzheimer’s disease (AD), especially in those at higher genetic risk (2) b) quercetin reduced AD like pathology in 3xTg-AD AD model mice (10) and c) PTPRD displayed robust genetic associations with densities of AD neurofibrillary pathology in AD brains (11).

We have found that a) 3xTg-AD/PTPRD+/- mice display hippocampal tau pathology at 4 months of age, b) structure activity relationship studies of more than 100 quercetin analogs identify 6BrQ as a superior PTPRD PAM, c) 3xTg-AD/PTPRD+/-mice are validated by their reduced pathological tau in mice treated with quercetin and d) 6BrQ in validated *in vivo via* much greater reductions in pathological tau in 3xTg-AD/PTPRD +/- mice than those provided by quercetin (5).

PTPRD is largely expressed in neurons in adult brain, providing cell-type specificity for modification of GSK3 activities in hyperphosphorylating tau and in contributing to neurofibrillary pathophysiology in AD (12). Our current observations that quercetin and 6BrQ both inhibit GSK3 directly add a possibly important, though less cell type specific, mechanism for the benefits that quercetin and other flavonols provide in mouse models and in humans. Quercetin has also been found to inhibit GSK3s in other work (13).

Quercetin and 6BrQ inhibition of cyclooxygenases 1 and 2 could also contribute to anti-inflammatory properties that could counter the inflammation that has been implicated in AD and other neurodegenerative processes (14) . Prior work noted quercetin inhibition of COX2 and modeled quercetin docking to COX1 and COX2 (15), though the primary data presented did not indicate inhibition (16).

Phosphodiesterase inhibition by quercetin has also been previously described, including some selectivity for PDE4 (17). Phosphodiesterase implication in models of cognitive decline and in models of neuroinflammation has led to interest in PDE inhibition as an anti-AD strategy (18).

Adenosine systems have also been linked to cognitive performance and to AD. EUROFINS notes that its assay monitors binding to SLC29A1, the equilibrative nucleoside transporter 1 (hENT1) that is a major player in purinergic signaling and is often responsible for adenosine uptake into nerve terminals (19). SLC29A1/ENT1 inhibitors can reverse AD-like mnemonic deficits in mouse models (20). Adenosine A2a receptors have also been strongly linked to AD pathophysiology (21). Findings from the current assays are supported by prior observations concerning quercetin interactions with adenosine receptors (22); interactions with SLC29A1 are less supported by previous work.

5HT2B serotonin receptor models can interact with quercetin *in silico* (23). Treatments with quercetin mimic benefits of drugs that impact 5HT2B receptors in blocking smoke-induced lung changes (24). Each of these observations is consistent with our current observations that demonstrate quercetin and 6BrQ blockade of these receptors at 10 μM concentrations. 5HT2B upregulation in AD brains and the involvement of this receptor in AD pathophysiology have both been reported (25).

Both quercetin and 6BrQ interact with the peroxisome proliferator-activated receptor gamma, a nuclear receptor/transcription factor, in the present data. These observations are consistent with prior reports that quercetin effects can be reversed *in vivo* by PPARgamma blockers (26).

Quercetin has attained FDA generally recognized as safe (GRAS) status and fails to display any well-documented toxicity at doses of several hundred milligrams/day in toxicologic studies (27). It is thus reassuring that 6BrQ displays only one activity in this large panel that is not displayed by quercetin. 6BrQ appears to gain potency at cholecystokinin 1 (CCK1) receptors that is not displayed by quercetin. CCK1 receptors are expressed in number of brain regions including nucleus tractus solitarius, area postrema, and hypothalamus as well as in the vagus nerve, gall bladder, pancreas and a number of other peripheral organs (28). Relatively selective CCK1 receptor antagonists, including dexloxiglumide, have been used in humans up to phase 3 trials without any unacceptable toxicity (29). There thus seems to be a low likelihood that 6BrQ’s modest potency at CCK1 receptors would impair its development toward human use.

By contrast, 6BrQ loses several properties displayed by quercetin, including inhibition of monoamine oxidase, activities at adenosine A1, and dopamine D4 G protein coupled receptors as well as interactions with progesterone and estrogen nuclear receptor/transcription factors. This difference in pattern of off target activities, combined with 6BrQ’s improved activity at PTPRD, provides an overall greater specificity for 6BrQ that for quercetin itself.

In conclusion, we report robust antioxidant effects of 6BrQ that are equivalent to those of quercetin itself. We provide the first comprehensive screen of quercetin effects on a large panel of targets of action of most known drugs, documenting modest potencies at only a few of these sites. We also provide the first off-target screening for 6BrQ, documenting modest potencies at an even smaller number of these sites that include several where 6BrQ effects could act along with those at PTPRD to reduce AD pathophysiology. Our observations add to the dynamic field of flavonoid pharmacology. They provide support for development of bioavailable glycosylated 6BrQ prodrugs to slow development of AD neurofibrillary pathology.

## Data availability

Underlying data is available from the authors on request.

## Competing interests

The US Department of Veterans Affairs has filed provisional patent applications that cover glycosylated bromoquercetins

## Ethics declaration

There is nothing to declare (note the above competing interest).

## Funding

We are grateful to support for this work from NIA (NIH, US) supplements to U01DA047713 and UG3DA056039 (GRU, PI) and to support from the Department of Neurology, University of Maryland School of Medicine and the VA Maryland Healthcare System.

## Author contributions

GRU Obtained funding, planned work, interpreted data, planned SAR and improved PTPRD PAM identification and wrote manuscript

BK Performed many *in vitro* orthophosphate assays for quercetin analog SAR.

IH Contributed to quercetin analog Structure-activity relationship studies and identification of 6BrQ including syntheses and initial orthophosphate assays. KS Contributed to quercetin analog SAR and 6BrQ identification, synthesis and purification.

## Acknowledgements

We are grateful to support from the grants noted above, with especial help from Drs David White, Matt Seager, Jane Acri, Nate Appel and Rik Kline (NIDA, NIH).

